# Cannabinoid during adolescence phenocopies reelin haploinsufficiency in prefrontal cortex synapses

**DOI:** 10.1101/2021.12.22.473793

**Authors:** T.J. Thenzing Juda Silva-Hurtado, Gabriele Giua, Olivier Lassalle, Michelle N. Murphy, Jim Wager-Miller, Ken Mackie, Olivier J. Manzoni, P. Pascale Chavis

## Abstract

In humans and rodents, the protracted development of the prefrontal cortex (PFC) throughout adolescence represents a time for marked vulnerability towards environmental adversities, such as stress or drug exposure. We previously showed that the extracellular matrix protein reelin is an instrumental synaptic modulator that shapes medial PFC’s (mPFC) circuitry during maturation and is a critical mediator of the vulnerability to environmental stress. Emerging evidence highlight the role of the endocannabinoid system in the postnatal maturation of the PFC and reelin deficiency influences behavioral abnormalities caused by heavy consumption of THC during adolescence. Could the reelin-dependent maturation of prefrontal networks may be vulnerable to cannabinoid exposure during adolescence? To explore this hypothesis, we studied the effects of a single *in-vivo* exposure to a synthetic cannabinoid on reelin expression and mPFC functions in adolescent male mice. The results show that a single cannabinoid exposure mimics reelin haploinsufficiency by decreasing prefrontal reelin expression in a layer-specific pattern without changing its transcriptional levels. Furthermore, this treatment impeded synaptic plasticity: adolescent cannabinoid lowered long-term potentiation to the magnitude observed in age-matched reelin haploinsufficient males. Quantitative PCR analysis showed that changes in the mRNA levels of NMDARs does not account for the reduction of TBS-LTP. Together, the data show that exposure to cannabinoid during adolescence phenocopies reelin haploinsufficiency and further identifies reelin as a key component of the vulnerability of PFC to environmental insults.

## INTRODUCTION

The prefrontal cortex (PFC) underlies multiple higher functions such as working memory, reasoning, cognitive flexibility, and emotionally guided behaviors. The PFC is one of the last structures of the central nervous system (CNS) to reach maturation. It continues to mature through early adulthood in humans and rodents (Gogtay et al. 2004; Chini and Hanganu-Opatz 2020). This protracted maturation is characterized by connectivity refinement and concomitant maturation of cognitive abilities in both humans and rodents (Giedd et al. 1996; Luna et al. 2001). Emerging evidence point to adolescence as a sensitive window during which external stimuli and experience shape changes in PFC structure and function but also a period for abnormal PFC maturation following exposure to environmental adversities, such as stress and drug exposure (Bava and Tapert 2010).

During adolescence the developing brain and in particular the PFC, is acutely sensitive to exogenous cannabinoids. Exogenous cannabinoid exposure during adolescence results in persistent PFC-dependent behavioral alterations and abnormalities in prefrontal architecture and synaptic functions (Levine et al. 2017; Miller et al. 2019; Renard et al. 2017; Zamberletti et al. 2014). The endocannabinoid system is comprised of the two G-protein coupled cannabinoid receptors (CB1 and CB2), as well as the endogenous cannabinoids ligands (anandamide and 2-arachidonoylglycerol) amongst other elements. The PFC is a brain area of dense expression of CB1 receptor (Marsicano and Lutz 1999; Lafourcade et al. 2007) and is consequently sensitive to various endocannabinoid-related synaptopathies which manifest as a variety of cognitive alterations (Scheyer et al 2017). Despite these advancements, the molecular mediators of the link between adolescent exposure to cannabinoids and prefrontal dysfunctions are unknown.

Reelin is a secreted glycoprotein of the extracellular matrix serving multiple functions in the brain throughout life. In the developing CNS reelin is essential to layer formation (Sekine et al. 2014) and in the postnatal and adult CNS it is an important contributor to the synaptic physiology (Jossin 2020). We previously showed that reelin is instrumental to the postnatal developmental trajectory of the prelimbic area of the medial prefrontal cortex (PL-PFC) at the structural, synaptic, and behavioral levels (Iafrati et al. 2014, 2016. Bouamrane et al. 2017). Our previous work showed that disruption of synaptic transmission and plasticity in the PL-PFC and of PFC-dependent behaviors by reelin haploinsufficiency occurs during adolescence (Iafrati et al. 2014, 2016. Bouamrane et al. 2017) and that reelin is an instrumental synaptic modulator underlying mPFC’s malfunctions in response to nutritional stress (Labouesse et al. 2017). Finally, reelin deficiency influences behavioral abnormalities caused by heavy consumption of THC during adolescence (Iemolo et al., 2021). Considering that reelin is critically involved in shaping adolescent PL-PFC circuitry and that adolescent PFC is highly vulnerable to disruption of the endocannabinoid system, it is intriguing to speculate that during adolescence the reelin-dependent maturation of prefrontal networks may be vulnerable to cannabinoid exposure. To address this question, we studied the effects of a single in vivo cannabinoid exposure in adolescent mice on reelin expression and synaptic plasticity in the PL-PFC.

## MATERIALS AND METHODS

### Animals and drug treatments

C57BL6/J males (Janvier-Labs, France) were received and left undisturbed during 7 days before drug administration. Drugs were injected intraperitoneally between postnatal days 34 and 40 (34-40), and mice were sacrificed 17±1 hours after drug administration. Animals received a single intraperitoneal injection of WIN55,212-2 (2mg/kg) alone or with SR141716A (8mg/kg). WIN55,212-2 or SR141716A were suspended in 1:2:37 dimethyl sulfoxide, cremophor, and NaCl 0.9% (B. Braun), and injected at 5 mL/kg. Control mice received vehicle. Mice were housed in standard 12h light–dark cycle and supplied food pellets and water *ad libitum*.

Experimenters were blind to treatments.

### Electrophysiology

Coronal slices containing the prelimbic area (PL) of the medial prefrontal cortex (mPFC) were prepared as previously described (Iafrati et al., 2014). Briefly, mice were anesthetized with isoflurane and 300 μm-thick coronal slices were prepared in a sucrose-based solution at 4°C using an Integraslice vibratome (Campden Instruments). Slices were stored for 30 min at 32°C in artificial cerebrospinal fluid (ACSF) containing (in mM): NaCl (130), KCl (2.5), MgCl_2_ (2.4), CaCl_2_ (1.2), NaHCO_3_ (23), NaH_2_PO_4_ (1.2) and Glucose (11), equilibrated with 95% O2 /5% CO2. Slices were then stored at room temperature until recording. All experiments were conducted at 30-32°C in ACSF. For recording, slices were superfused at 2 ml per min with ACSF containing Picrotoxin (100µM; Sigma) or SR95531 (Gabazine, 5µM; Tocris) to block GABA_A_ receptors.

The prelimbic area of the mPFC (PL-PFC) was visualized using an infrared illuminated upright microscope (Olympus BX51WI, France) and extracellular field excitatory postsynaptic potentials (fEPSPs) recordings carried out as previously described (Iafrati et al., 2014; 2016). fEPSPs were recorded in the PL-PFC layer V with an ACSF-filled electrode and evoked in layer III at 0.1 Hz with a stimulating glass electrode filled with ACSF. LTP was induced using a theta-burst stimulation protocol consisting of five trains of burst with four pulses at 100 Hz, at 200 ms interval, repeated four times at intervals of 10 s. The glutamatergic nature of fEPSPs was confirmed at the end of each experiment by perfusing the non-NMDA (N-methyl-D-aspartate) ionotropic glutamate receptor antagonist 6-cyano-7-nitroquinoxaline-2,3-dione (CNQX) (20 μM; NIH), which specifically blocked the synaptic component without altering the non-synaptic component (not shown)

Signals were collected using an Axopatch-200B amplifier (Axon Instruments, Molecular Devices, Sunnyvale, USA), filtered at 2 kHz, digitized at 10 kHz, acquired with Clampex 10.7 acquisition Software via a Digidata 1440A (Axon Instruments) and analyzed using Axograph 1.7.6.

### Immunohistochemistry

Animals were deeply anesthetized with Pentobarbital (90mg/kg; Exagon Med’Vet) and perfused transcardially with cold phosphate-buffered saline solution (PBS; Gibco Life Technologies) followed with Antigenfix (DiaPath) a phosphate-buffered paraformaldehyde solution. The dissected brains were postfixed overnight at 4°C in the same fixative. Brains were then sectioned using a vibratome (VT 1200 s, Leica) into 60 μm-thick coronal slices.

Sections were first rinsed three times for 10 min in PBS and then incubated in blocking solution with 0.1M PBS containing 0.3% Triton X100 (Sigma) and 0.2% Bovine Serum Albumin (BSA; Sigma) twice for 1h at room temperature (RT). Slices were incubated free-floating overnight at RT with the mouse G10 anti-reelin primary antibody (1:3000, MAB5406; Millipore) diluted in the blocking solution. After three blocking solution washes (10 min each), section was incubated at RT for 75 min with the secondary antibody donkey anti-mouse Alexa 568 (Invitrogen ThermoFisher Scientific) diluted 1:500 in the blocking solution. After three rinses of 10 min in 0.1 M PBS, sections were stained with Hoechst (Invitrogen ThermoFisher Scientific) diluted 1:1000 in 0.1M PBS for 12 min, washed again three times for 10 min in 0.1 M PBS and coverslipped with Aqua-Poly/Mount (Polysciences).

The specificity of the G10 antibody was tested on sections obtained from homozygous reeler mice lacking reelin expression. Additional negative controls were performed by omitting the G10 primary antibody on wild-type or HRM sections.

### Image analysis

Confocal images were acquired with a Zeiss LSM-800 system equipped with emission spectral detection and a tunable laser providing excitation range from 470 to 670 nm. Stacks of optical sections were collected with a Plan Apochromat 20x.0.8 air objective for 3D reconstructions. Laser power and photomultiplier gain were adjusted to obtain few pixels with maximum intensity on the somata showing the higher labelling intensity. To obtain a whole rostro-caudal representation of the PL-PFC of each mouse, images were acquired from Bregma 2.58 to 1.94 according to the Mouse Brain Atlas (Paxinos & Franklin 2009). To obtain a Z-representation across layers I to VI within each brain section, scanning was performed using tiles representing a total of 894.4 μm x 894.4 μm surface/image size and a Z-stack selection covering a depth of 17.5 to 19.8 µm. The tri-dimensional reconstruction and blind semiautomated analysis were performed with Fiji (Image J).

### Western blots

Immunoblotting of reelin in PFC lysates were performed following published procedures (Sinagra et al. 2005).

### Quantitative Reverse Transcriptase PCR

Brains were harvested and snap frozen in isobutane on dry ice and stored at -80°C. Frozen rat brains were coronally sectioned in a cutting block (Braintree Scientific, Inc., Braintree, MA,) that had been pre-chilled to -20°C. One-millimeter sections were kept frozen throughout dissection with brain regions stored at -80°C until use. Quantitative reverse transcriptase PCR was performed on mPFC. Total RNA was extracted using the RNeasy Plus Micro Kit (Qiagen, Hilden, Germany). RNA was reverse transcribed using the RevertAid Kit (Thermo Fisher Scientific, Waltham, MA) as per manufacturer’s instructions. All Taqman probe sets used were from Thermo Fisher Scientific, USA: RELN (Mm00465200_m1), GRIN1(Mm00433790_m1), GRIN2A (Mm00433802_m1), GRIN2B (Mm00433820_m1) and GRIN2D (Mm00433822_m1). TaqMan Gene Expression Master Mix from Applied Biosystems (Foster City, CA) was used in generating expression data on a QuantStudio7 thermal cycler. Duplicates were run for each sample, and relative gene expression was determined using the double delta Ct method.

### Statistical Analysis

All values are given as mean ± SEM and statistical significance was set at *P* < 0.05. Statistical analysis was performed with GraphPad Prism 9.2.0 (GraphPad Software, La Jolla, CA, USA). Multiple comparisons were made using a one-way analysis of variance (ANOVA) followed, if significant, by Tukey’s multiple comparisons test. Two sample comparisons were made with two-tailed paired t-tests.

## RESULTS

### Adolescent exposure to a synthetic cannabimimetic alters prefrontal reelin expression in a layer specific manner

Adolescent male mice received a single intraperitoneal injection of the synthetic cannabinoid WIN55,212-2 and the following day reelin expression pattern was studied in the PL-PFC using immunofluorescence.

We found that WIN exposure reduced the density of reelin-positive cells by 18.2 ±1.6% in the whole PL-PFC compared to naïve and vehicle-treated mice (Figure 1A). We analyzed whether this effect was similar across all PL-PFC layers. In layer I, the cumulative distributions of reelin-positive cell densities obtained in naïve, vehicle- and WIN-treated mice were superposable, showing a lack of WIN effect in the most superficial layer of the PL-PFC (Figure 1B). In contrast, in layers II/III and V/VI, the distribution of the WIN-treated group was shifted to the left compared to naïve and vehicle-treated mice, indicating that WIN exposure decreased the density of reelin-positive cells in these layers (Figure 1B). Coadministration of the CB1 receptor antagonist SR141716A with WIN55,212-2 prevented the reduction in the total density of reelin-positive cells (Figure 1A). It also normalized the densities distributions in layers II/III and V/VI as shown by the overlap of SR+WIN, vehicle, and naive cumulative curves (Figure 1B). Taken together, these observations show that the effect of the cannabinoid WIN55,212-2 is layer specific and mediated by CB1 receptors activation.

**Figure 1:**
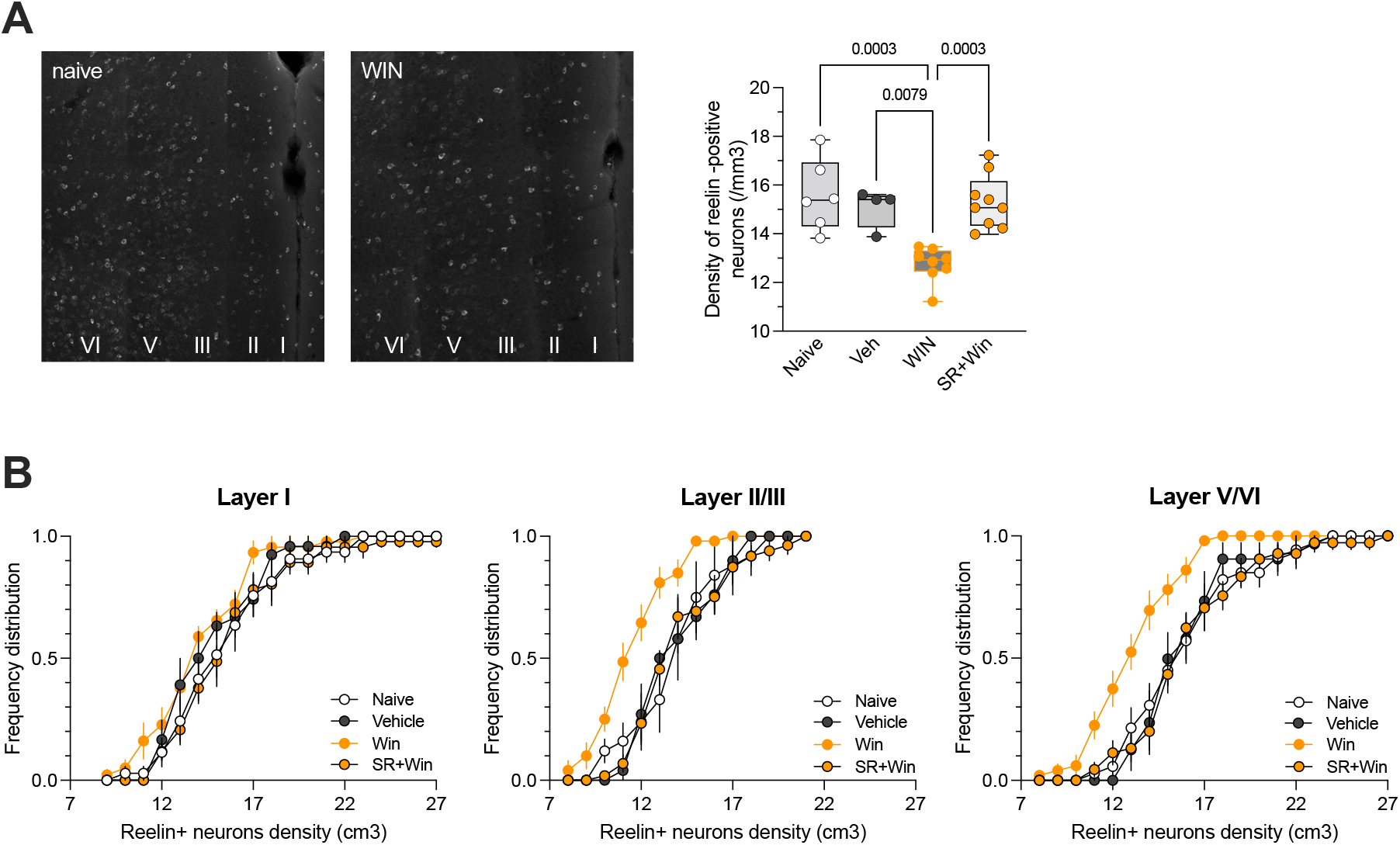
A single in vivo exposure to cannabinoid decreases the density of reelin-expressing cells in a layer specific manner. **A**: Confocal images of PL-PFC sections stained for reelin from P40 naive and WIN-treated mice. Box and whiskers plot showing median, 25^th^ to 75^th^ percentiles, minimum to maximum of the density of reelin-positive cells per volume as well as individual values obtained for each mouse. The median was 15.4 in naive mice (n=6), 15.4 in vehicle-treated mice (n=4), 12.9 in WIN-treated mice (n=8) and 15.1 in SR+WIN-treated mice (n=9). F(_3,23_)=11.44, P<0.0001, ANOVA. Only P values >0.05 are displayed on the graph. **B:** The cumulative distributions of the densities of reelin-positive cells across the different PL-PFC layers show a specific effect of WIN in layers II/III and layers V/VI. This effect is prevented by SR141716A. Values are means ± sem.

### Cannabinoid exposure downregulates reelin expression without altering transcriptional levels

We next examined whether the WIN-induced reduction of prefrontal reelin-positive cells density was correlated with expression changes in reelin levels. Western blot analyses were performed on PFC of vehicle-, WIN- and SR+WIN-exposed mice (Figure 2A-B). We found a significant decrease of the total reelin expression in WIN-injected mice that was blocked by the CB1 receptors antagonist SR141716A, indicating that CB1 receptor activation reduces the total amount of reelin in the PL-PFC (Figure 2A-B). In contrast, CB1 knock-out mice showed no reduction in total reelin levels when injected with WIN compared to WIN-treated wild-type littermates (Figure 2C). Altogether, these results show that WIN exposure reduces reelin expression through CB1R activation.

**Figure 2:**
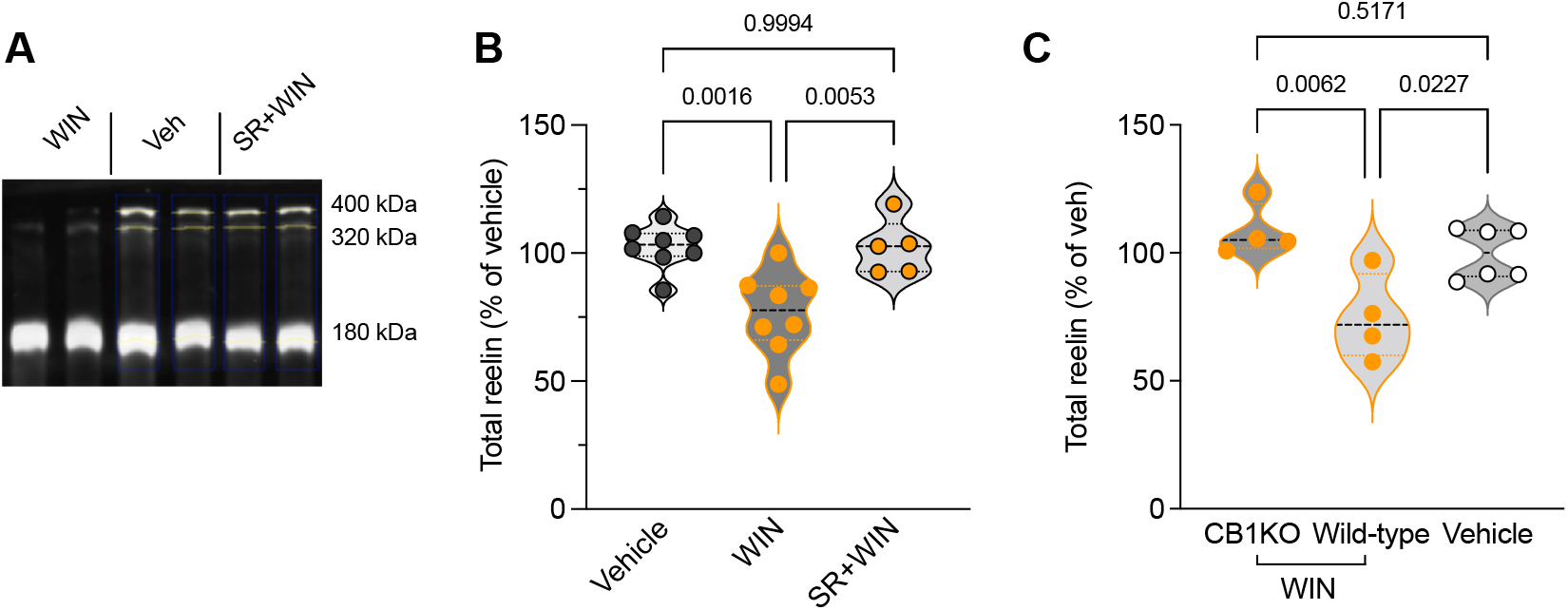
CB1 receptor activation decreases prefrontal reelin expression. **A:** Representative immunoblotting of whole PFC lysates extracted from vehicle-, WIN- and SR+WIN-exposed mice and probed with the anti-reelin G10. Each lane is a different mouse. **B:** Densitometric measurements of total reelin levels (sum of the 3 isoforms) expressed as the percentage of total reelin levels in vehicle treated mice. Violin plot shows that WIN 55,212-2 decreased reelin levels (n=8) compared to vehicle treated mice (n=8) and that his effect is prevented by the antagonist SR141716A (n=5). F_(2,18)_ =10.65, P=0.0009, ANOVA. **C:** Densitometric analysis of total reelin levels after WIN injection in CB1KO mice (n=4) and their wild-type littermates (n=4) compared to vehicle-injected CB1KO and vehicle-injected wild-type littermates (Vehicle, n=6). Reelin levels were not reduced by WIN administration in CB1KO compared to wild-type littermates. F_(2,11)_=8.406, P=0.006, ANOVA. n is the number of mice per group.

In the developing and adult brain, after secretion the full-length reelin protein (relative molecular mass of 388 kDa, and 450kDa when glycosylated; D’Arcangelo et al. 1995) is enzymatically cleaved at two major sites (Jossin et al. 2007; Krstic et al. 2012). This process generates five fragments, of which two are detected by the G10 antibody recognizing epitopes in the N-terminal region (de Bergeyck et al., 1998; Sinagra et al. 2005; Groc et al. 2007). When analyzing the different forms recognized by the antibody, we observed a decrease in the amount of full-length protein and the 320kDa cleavage product in the WIN-exposed group (Supplementary Figure 1). These results suggest that secretion is downregulated, and that cleavage may be impacted by WIN exposure.

To gain more mechanistic insight into reelin downregulation in cannabinoid-exposed mice, we performed quantitative PCR on the medial PFC from naive, vehicle- or WIN-exposed mice (Figure 3A). No difference in reelin mRNA levels was found between the different groups (Figure 3A), showing that adolescent cannabinoid exposure does not affect reelin transcription.

**Figure 3:**
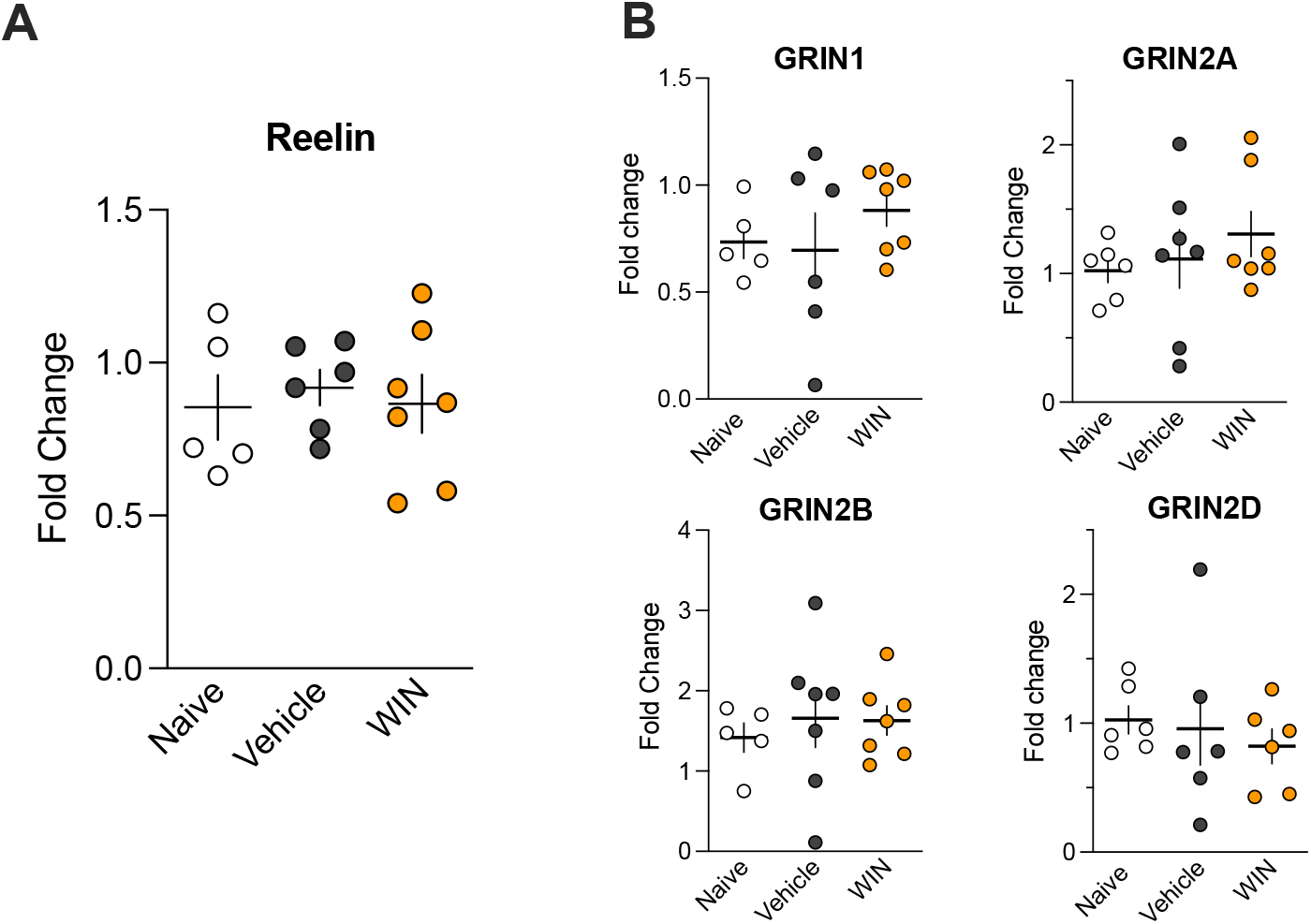
Adolescent Win 55,212-2 exposure alters neither reelin nor NMDAR subunits messenger RNA. **A:** Quantitative polymerase chain reaction analysis of reelin mRNA in medial PFC tissue collected from naive, vehicle and WIN55,212-2-treated mice. F_(2, 15)_= 0.1450, P=0.8662. **B:** Messenger RNA levels of GluN1 (F_(2, 15)_ = 0.7564, P=0.4865), GluN2A (F_(2, 17)_ = 0.6354, P=0.5418), GluN2B (F_(2, 16)_=0.2040, P=0.8175) and GluN2D (F_(2, 15)_= 0.3019, P=0.7438) measured in the same individuals as in A. All values are means ± sem and each circle represents an individual animal.

### Adolescent cannabinoid exposure reduces NMDAR-dependent plasticity at layer V/VI excitatory synapses in PL-PFC

We previously reported that reduced levels of reelin alter deep layers excitatory synaptic plasticity in the PL-PFC during adolescence (Iafrati et al. 2016). Considering that CB1 receptor activation downregulates reelin expression in layers II/III and V/VI, we examined the effect of a single WIN injection on synaptic plasticity of deep layers excitatory synapses. We used a theta-burst stimulation protocol in layer III to induce long-lasting synaptic potentiation (TBS-LTP) in layer V. This protocol induced a robust LTP at layer III/V excitatory synapses in naïve (40.8 ± 4.0%, n=7) and vehicle-treated mice (40.0 ± 5.3%, n=6; Figure 4). In contrast, the LTP magnitude was markedly reduced in WIN-treated mice (14.5 ± 3.0%, n=9). Following SR+WIN exposure, it was normalized to the levels of naive- and vehicle-treated mice (34.5 ± 4.7%, n=7; Figure 4). Altogether, these data show that activation of CB1 receptors reduces TBS-LTP at deep PL-PFC excitatory synapses. This effect could not be attributed to differences in the excitability of layer III/V excitatory synapses as input-output curves were identical between the different groups (Supplementary Figure 2).

**Figure 4:**
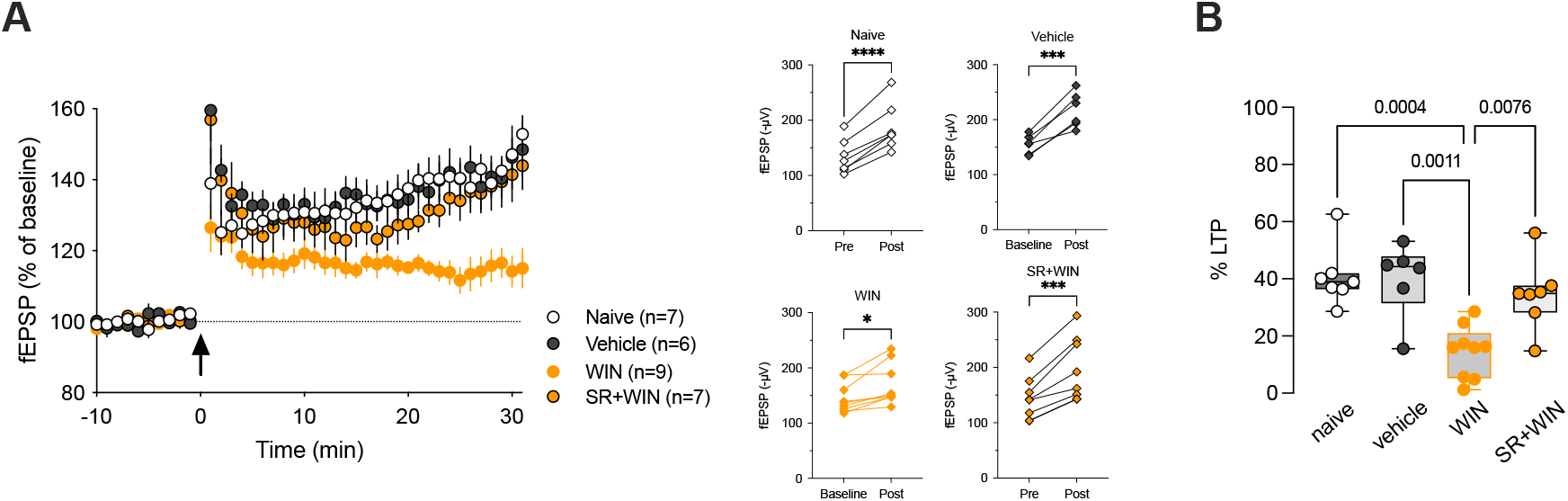
Disruption of NMDAR-dependent long-term potentiation at layer V/VI excitatory synapses in PL-PFC after a single in vivo cannabinoid exposure. **A:** Grouped time courses of field excitatory postsynaptic potential (fEPSP) responses expressed as the percentage of baseline before and after theta-burst stimulation (TBS, indicated by arrow). TBS induced LTP with similar time courses in naive, vehicle- and SR141716A+WIN55,212-2-treated mice. TBS-LTP was reduced in WIN55,212-2 exposed mice. Right: fEPSPs amplitudes of individual experiments before (baseline) and 20-30 min after TBS (post). **B:** Box and whiskers plot showing individual values, median, 25^th^ to 75^th^ percentiles, minimum to maximum of the percentage of potentiation measured 20-30 min after TBS. F_(3,25)_=9.968, P=0.0002, ANOVA. Only P values >0.05 are displayed on the graph. n represents the number of individual mice.

Considering that this form of plasticity is *N*-methyl-D-aspartate receptors (NMDAR)-dependent (Iafrati et al. 2014), we examined whether the transcriptional levels of the different NMDAR subunits were affected following WIN-exposure. Quantitative PCR analysis did not show changes in the mRNA levels of NMDARs subunits in the medial PFC of WIN-treated mice compared to naïve and vehicle groups (Figure 3B). These data indicate that the reduction of TBS-LTP in WIN-exposed mice does not result from changes in the transcription of NMDAR subunits.

To uncover a possible relationship between reelin deficiency and reduced TBS-LTP following WIN administration, we examined TBS-induced plasticity in a mutant mouse model haploinsufficient for reelin (Iafrati et al. 2016). Interestingly, we observed that the time course and the magnitude of TBS-LTP in adolescent heterozygote reeler male mice (HRM) are like those observed in WIN-exposed animals (14.5 ± 3.0%, n=9 WIN-treated and 13.1± 3.7%, n=9 HRM; Supplementary Figure 3). These results show that constitutive decreased levels of endogenous reelin mimics the effect of WIN-induced downregulation of reelin expression on the plasticity of layer III/V excitatory synapses of the PL-PFC. This observation raises the hypothesis that alteration of TBS-induced plasticity following in vivo exposure to the cannabimimetic WIN55,212-2 is a consequence of the decrease in endogenous reelin levels triggered by WIN exposure.

## DISCUSSION

We found that a single in vivo exposure of adolescent male mice to the cannabimimetic WIN55,212-2 decreases the density of reelin-positive neurons in the PL-PFC in a layer specific manner. This effect resulted from the downregulation of prefrontal reelin protein expression without changes in reelin transcriptional levels. The WIN-55,212-2-mediated effects were prevented by administration of a CB1 receptor antagonist, thus confirming mediation of synthetic cannabinoid exposure effects by CB1 receptor. Finally, WIN55,212-2 exposure phenocopied reelin haploinsufficiency and disrupted TBS-induced long-term potentiation at excitatory synapses between layers III and V. Altogether, these results point towards a vulnerability of adolescent PL-PFC to cannabinoid exposure mediated by a deficit in reelin protein induced by CB1 receptor activation.

In rats, a single exposure to synthetic CB1 receptor agonist WIN55,212-2 for 24 hours has been reported to abolish TBS-LTP in adults but not at the juvenile stage (Borsoi et al. 2019). In rats, TBS-LTP ablation is also sex specific as it was not observed in females (Borsoi et al. 2019). TBS-LTP relies on NMDAR activation and evidence indicates that NMDAR activity may be differently modulated according to sex (McRoberts et al. 2007). Moreover, it was reported that hormonal status influences both long-term potentiation induction and NMDAR function in male rats (Moradpour et al., 2013). Considering this evidence, it may be relevant to further investigate whether adolescent exposure to WIN also affect TBS-LTP in female mice.

A limitation of our study is that our experiments did not ascertain that the reduction of TBS-LTP following WIN exposure resulted from a decrease in reelin expression. We show that both the magnitude and the time course of synaptic plasticity in WIN-treated mice and HRM are indistinguishable (Supplementary Figure 2). To resolve this limitation, recombinant reelin will be injected directly into the medial PFC according to our previous work (Labouesse et al. 2017; Rogers et al. 2011), several days before WIN-exposure and assessment of long-term synaptic plasticity.

Our study revealed for the first time that the activation of the endocannabinoid system through CB1 receptors downregulates reelin expression in the PL-PFC in a layer-specific manner. Importantly, in the PL-PFC, CB1 receptor expression pattern was shown to be restricted to layers II/III and V/V (Lafourcade et al. 2007). This selective expression pattern provides an interpretation for the lack of reduction in the density of reelin-expressing cells by WIN55,212-2 in layer I.

Whether a single exposure to WIN has long lasting effects on reelin expression and whether chronic cannabinoid exposure could further decrease reelin levels in wild-type mice remain to be tested. A recent study reported that reelin haploinsufficiency favors the vulnerability to adolescent chronic exposure to the psychoactive component of cannabis, Δ9-tetrahydrocannabinol (THC; Iemolo et al. 2021). Chronic exposure to high doses of THC during adolescence caused pronounced behavioral alterations in adults HRM in a sex-specific manner (Iemolo et al. 2021). This further supports the hypothesis that WIN-induced reelin deficits participate to the functional and behavioral impairments triggered in the PFC following adolescent cannabinoid exposure (Renard et al. 2014; Rubino et Parolaro. 2016). Hence, the molecular link between CB1 receptor and reelin signaling remains to be elucidated.

Finally, we recently reported that reelin is a sensitive target to adolescent high fat exposure and a molecular mediator of the link between adolescent high fat feeding stress and adult deficits in prefrontal cognitive functions (Labouesse et al. 2017). Collectively this study and the present data, demonstrate that reelin is a sensitive target of various early-life environmental insults, such as stress and drug exposure and may be the neurobiological hub underlying the development of psychiatric diseases.

## Author contributions

Thenzing Juda Silva Hurtado: Conceptualization; Data curation; Formal analysis; Validation.

Gabriele Giua: Data curation; Formal analysis; Validation; Methodology.

Olivier Lassalle: Data curation; Validation; Methodology.

Michelle Murphy: Data curation; Formal analysis; Validation; Methodology.

Jim Wager-Miller: Data curation; Formal analysis; Validation; Methodology.

Ken Mackie: Conceptualization; Methodology and editing.

Olivier JJ Manzoni: Conceptualization; Methodology and editing.

Pascale Chavis: Conceptualization; Supervision; Funding acquisition; Data curation;

Formal analysis; Validation; Methodology; Writing: original draft, review and editing;

Project administration.

## Acknowledgements

The authors are grateful to D. Verrier for the western blots and to Dr. G. Marsicano for the gift of the CB1R knock-out mice. This work was supported by the Institut National de la Santé et de la Recherche Médicale (INSERM), Agence Nationale de la Recherche (ANR Cannado) and the NIH (5R01DA043982-02). The authors are grateful to the Chavis-Manzoni team members for helpful discussions. We thank the National Institute of Mental Health’s Chemical Synthesis and Drug Supply Program (Rockville, MD, USA) for providing CNQX, WIN 212,212,2 and and SR141716A.

## Declarations of interest

The authors declare no competing interests.

**Supplementary Figure 1:**
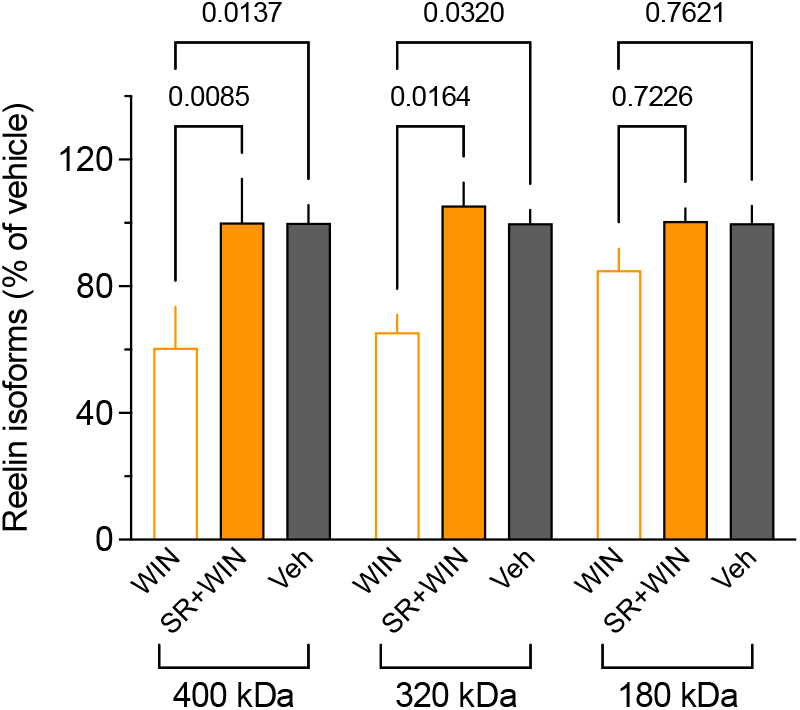
Effect of WIN exposure on reelin cleavage fragments. Densitometric analysis of full-length reelin (400kDa), and the two cleaved products (320 and 180 kDa) detected by the G10 antibody in WIN-(n=8), SR+WIN-(n=5) and vehicle (n=8). Values are expressed as the percentage of the corresponding reelin isoform detected in the vehicle group. F_(8,54)_=4.095, P=0.0012, ANOVA. Values are means ± sem and n is the number of mice per group.

**Supplementary Figure 2:**
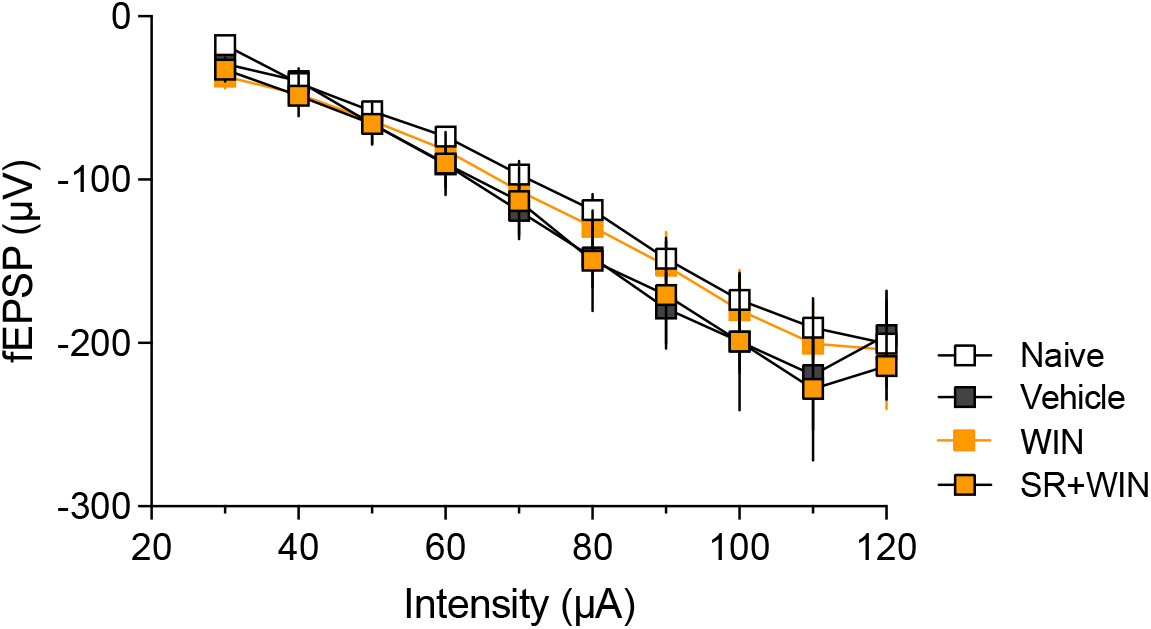
Excitability of layer III/V synapses is not affected by the different treatments. Input-output profile of average fEPSP amplitude (± sem) is shown for the different groups of mice: naïve (n=7), vehicle (n=6), WIN55,212-2 (WIN, n=9) and SR141716A + WIN55,212-2 (SR+WIN, n=7).

**Supplementary Figure 3:**
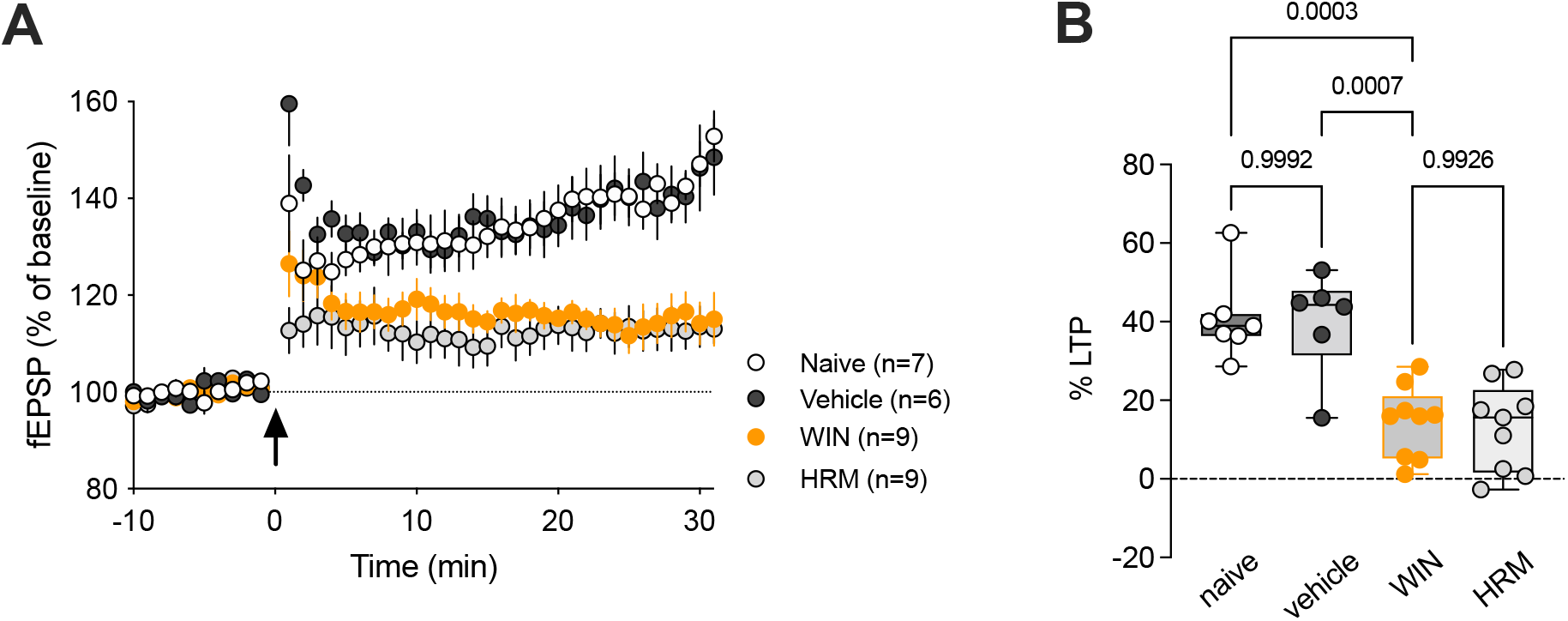
Adolescent Win 55,212-2 exposure phenocopies reelin haploinsufficiency. **A:** Time course of TBS-LTP normalized to baseline in adolescent HRM, naive, vehicle- and WIN55,212-2-treated male mice. **B:** Box and whiskers plot showing individual values, median, 25^th^ to 75^th^ percentiles, minimum to maximum of the percentage of potentiation measured 20-30 min after TBS. F_(3,27)_=15.27, P<0.0001, ANOVA. n represents individual animals.

## Notes

### Competing Interest Statement

The authors have declared no competing interest.

## REFERENCES

Bava S. & Tapert S.F. Adolescent Brain Development and the Risk for Alcohol and Other Drug Problems. Neuropsychol Rev (2010) 20:398–413.

Borsoi M., Manduca A. Bara A., Lassalle O., Pelissier-Alicot A.L. and Manzoni O.J.J. Sex Differences in the Behavioral and Synaptic Consequences of a Single in vivo Exposure to the Synthetic Cannabimimetic WIN55,212-2 at Puberty and Adulthood. Front. Behav. Neurosci., 2019 Mar 5;13:23

Bouamrane L, Scheyer AF, Lassalle O, Iafrati J, Thomazeau A, Chavis P. Reelin-Haploinsufficiency Disrupts the Developmental Trajectory of the E/I Balance in the Prefrontal Cortex. Front Cell Neurosci. 2017 Jan 12;10:308. doi: 10.3389/fncel.2016.00308. eCollection 2016.

Chini M, Hanganu-Opatz IL. Prefrontal Cortex Development in Health and Disease: Lessons from Rodents and Humans. Trends Neurosci. 2021 Mar;44(3):227–240. doi: 10.1016/j.tins.2020.10.017. Epub 2020 Nov 24.

D’Arcangelo, G.; Miao, G.G.; Chen, S.-C.; Scares, H.D.; Morgan, J.I.; Curran, T. A protein related to extracellular matrix proteins deleted in the mouse mutant reeler. Nature 1995, 374, 719–723.

de Bergeyck, V., Naerhuyzen, B., Goffinet, A.M., and Lambert de Rouvroit, C. (1998). A panel of monoclonal antibodies against reelin, the extracellular matrix protein defective in reeler mutant mice. J Neurosci Methods 82, 17–24.

Giedd J. N., A C Vaituzis, S D Hamburger, N Lange, J C Rajapakse, D Kaysen, Y C Vauss, J L Rapoport. Quantitative MRI of the temporal lobe, amygdala, and hippocampus in normal human development: ages 4-18 years. J Comp Neurol 1996 Mar 4;366(2):223–30.

Gogtay, N., Giedd, J. N., Lusk, L., Hayashi, K. M., Greenstein, D., Vaituzis, A. C., et al. (2004). Dynamic mapping of human cortical development during childhood through early adulthood. Proc. Natl. Acad. Sci. U.S.A. 101, 8174–8179. doi: 10.1073/pnas.0402680101

Iafrati, J., Malvache, A., Gonzalez Campo, C., Orejarena, M. C., Lassalle, O., Bouamrane, O., et al. (2016). Multivariate synaptic and behavioral profiling reveals new developmental endophenotypes in the prefrontal cortex. Sci. Rep. 6, 35504. doi: 10.1038/srep35504

Iafrati, J., Orejarena, M. J., Lassalle, O., Bouamrane, L., Gonzalez-Campo, C., and Chavis, P. (2014). Reelin, an extracellular matrix protein linked to early onset psychiatric diseases, drives postnatal development of the prefrontal cortex via GluN2B-NMDARs and the mTOR pathway. Mol. Psychiatry 19, 417–426. doi: 10.1038/mp.2013.66

Iemolo A., Montilla-Perez P., Nguyen J., Victoria B., Risbrough V.B., Taffe M.A., Francesca Telese F. Reelin deficiency contributes to long-term behavioral abnormalities induced by chronic adolescent exposure to Δ9-tetrahydrocannabinol in mice. Neuropharmacology. 2021 Apr 1;187:108495. doi: 10.1016/j.neuropharm.2021.108495. Epub 2021 Feb 11.

Jossin, Y.; Gui, L.; Goffinet, A.M. Processing of Reelin by Embryonic Neurons Is Important for Function in Tissue But Not in Dissociated Cultured Neurons. J. Neurosci. 2007, 27, 4243–4252

Jossin Y. Reelin Functions, Mechanisms of Action and Signaling Pathways During Brain Development and Maturation. Biomolecules. 2020 Jun 26;10(6):964. doi: 10.3390/biom10060964.

Krstic D., Rodríguez M., Knuesel I. Regulated Proteolytic Processing of Reelin through Interplay of Tissue Plasminogen Activator (tPA), ADAMTS-4, ADAMTS-5, and Their Modulators. PLoS ONE 2012, 7, e47793.

Katsutoshi S., Kubo K., Nakajima K. How does Reelin control neuronal migration and layer formation in the developing mammalian neocortex? Neurosci Res. 2014 Sep;86:50–8. doi: 10.1016/j.neures.2014.06.004. Epub 2014 Jun 23

Hypervulnerability of the adolescent prefrontal cortex to nutritional stress via reelin deficiency M A Labouesse, O Lassalle, J Richetto, J Iafrati, U Weber-Stadlbauer, T Notter, T Gschwind, L Pujadas, E Soriano, A C Reichelt, C Labouesse, W Langhans, P Chavis & U Meyer. Hypervulnerability of the adolescent prefrontal cortex to nutritional stress via reelin deficiency. Molecular Psychiatry volume 22, pages 961–971 (2017)

Lafourcade M., IElezgarai I., Mato S., Bakiri Y., Grandes, P. Manzoni OJJ. Molecular components and functions of the endocannabinoid system in mouse prefrontal cortex. PLoS One 2007 Aug 8;2(8):e709. doi: 10.1371/journal.pone.0000709.

Levine A., Clemenza K., Rynn M., Lieberman J. Evidence for the Risks and Consequences of Adolescent Cannabis Exposure. Am Acad Child Adolesc Psychiatry. 2017 Mar;56(3):214–225. doi: 10.1016/j.jaac.2016.12.014.

Luna B, Thulborn KR, Munoz DP, Merriam EP, Garver KE, Minshew NJ, Keshavan MS, Genovese CR, Eddy WF, Sweeney JA. Maturation of widely distributed brain function subserves cognitive development. Neuroimage. 2001 May;13(5):786–93. doi: 10.1006/nimg.2000.0743.

Marsicano G, Lutz B. Expression of the cannabinoid receptor CB1 in distinct neuronal subpopulations in the adult mouse forebrain. Eur J Neurosci. 1999;11:4213–25.

McRoberts J.A., Li J., Ennes H S, Mayer E A. Sex-dependent differences in the activity and modulation of N-methyl-d-aspartic acid receptors in rat dorsal root ganglia neurons Neuroscience. 2007 Sep 21;148(4):1015–20.

Miller ML, Chadwick B, Dickstein DL, Purushothaman I, Egervari G, Rahman T, Tessereau C, Hof PR, Roussos P, Shen L, Baxter MG, Hurd YL. Adolescent exposure to Δ9-tetrahydrocannabinol alters the transcriptional trajectory and dendritic architecture of prefrontal pyramidal neurons. Mol Psychiatry. 2019 Apr;24(4):588–600. doi: 10.1038/s41380-018-0243-x. Epub 2018 Oct 3.

Moradpour, F., Fathollahi, Y., Naghdi, N., Hosseinmardi, N., and Javan, M. (2013). Prepubertal castration causes the age-dependent changes in hippocampal long-term potentiation. Synapse 67, 235–244. doi: 10.1002/syn.21636

Renard J, Rosen LG, Loureiro M, De Oliveira C, Schmid S, Rushlow WJ, Laviolette SR. Adolescent Cannabinoid Exposure Induces a Persistent Sub-Cortical Hyper-Dopaminergic State and Associated Molecular Adaptations in the Prefrontal Cortex. Cereb Cortex. 2017 Feb 1; 27(2):1297–1310.

Renard, J., Krebs, M.O., Le Pen, G., Jay, T.M., 2014. Long-term consequences of adolescent cannabinoid exposure in adult psychopathology. Front. Neurosci. 8, 361.

Rogers JT, Rusiana I, Trotter J, Zhao L, Donaldson E, Pak DT, Babus LW, Peters M, Banko JL, Chavis P, Rebeck GW, Hoe HS, Weeber EJ. Reelin supplementation enhances cognitive ability, synaptic plasticity, and dendritic spine density. Learn Mem. 2011 Aug 18;18(9):558–64. doi: 10.1101/lm.2153511. Print 2011 Sep.

Rubino, T., Parolaro, D., 2016. The impact of exposure to cannabinoids in adolescence: insights from animal models. Biol. Psychiatr. 79, 578–585.

Scheyer AF, Martin HGS, Manzoni OJ. The endocannabinoid system in prefrontal synaptopathies. In: Melis M. editor Endocannabinoids and Lipid Mediators in Brain Functions. Springer, Cham; 2017.

Sinagra, M., Verrier, D., Frankova, D., Korwek, K. M., Blahos, J., Weeber, E. J., et al. (2005). Reelin, very-low-density lipoprotein receptor, and apolipoprotein E receptor 2 control somatic NMDA receptor composition during hippocampal maturation in vitro. J. Neurosci. 25, 6127–6136. doi: 10.1523/JNEUROSCI.1757-05.2005

Zamberletti E, Beggiato S, Steardo L, Prini P, Antonelli T, Ferraro L, Rubino T, Parolaro D (2014) Neurobiology of Disease Alterations of prefrontal cortex GABAergic transmission in the complex psychotic-like phenotype induced by adolescent delta-9-tetrahydrocannabinol exposure in rats. Neurobiol Dis 63:35–47. 10.1016/j.nbd.2013.10.028

